# Multiscale radiobiological assessment of laser-driven very high energy electrons versus conventional electrons

**DOI:** 10.1101/2025.05.27.656200

**Authors:** Camilla Giaccaglia, Emilie Bayart, Maxime Dubail, Chaitanya Varma, Sophie Heinrich, Julien Gautier, Amar Tafzi, Olena Kononenko, Jean-Philippe Goddet, Isabelle Lamarre-Jouenne, Charles Fouillade, Alessandro Flacco

**Affiliations:** Laboratoire d’Optique Appliquée, ENSTA Paris, CNRS, Ecole Polytechnique, Institut Polytechnique de Paris, 91120 Palaiseau, France; Institut Curie, Inserm U1021-CNRS UMR 3347, Paris Saclay University, Centre Universitaire, 91405 Orsay Cedex, France; Laboratory for Optics and Biosciences, Ecole polytechnique, CNRS, INSERM, Paris Saclay University, 91128 Palaiseau Cedex, France

**Keywords:** Very High Energy Electrons (VHEE), Conventional Intermediate Energy Electrons (CIEE), Laser-plasma accelerator (LPA), Linear accelerator (LINAC), Radiation-induced toxicity, Cell survival, Precision-cut lung slices (PCLS), Zebrafish embryos

## Abstract

**Purpose:** This study systematically investigates the radiobiological effects of Very High Energy Electrons (VHEE) generated by a laser-plasma accelerator (LPA), in comparison with Conventional Intermediate Energy Electrons (CIEE) from a conventional linear accelerator (LINAC). Using *in vitro, ex vivo*, and *in vivo* models, we evaluate and compare their potential toxicity on healthy tissues.

**Methods and Materials:** Cell survival, tissue response, and developmental toxicity were assessed across three biological models. *In vitro*, human fibroblasts (MRC5-hTERT) were used to generate post-irradiation survival curves. *Ex vivo*, precision-cut lung slices (PCLS) from mice were analyzed for radiation-induced inhibition of cell proliferation. *In vivo*, zebrafish embryos were used to evaluate developmental toxicity through body length and spinal curvature measurements. VHEE irradiations were performed using a broadband electron beam spanning 50–300 MeV, using the Salle Jaune LPA (Laboratoire d’Optique Appliquée, France), while CIEE exposures were performed with a 7 MeV conventional LINAC (Institut Curie, France).

**Results:** *In vitro*, MRC5-hTERT cells showed no significant difference in radiosensitivity between VHEE and CIEE, with comparable D_10_ values (p-value = 0.7). In the *ex vivo* model, both beams induced a dose-dependent decrease in cell division with no significant inter-beam differences at any dose level (p-value > 0.99). *In vivo*, zebrafish embryos exhibited dose-dependent body shortening and increased spinal curvature following both VHEE and CIEE exposure. No significant differences were observed between the two modalities at matched doses for any measured metric (p-value *≥* 0.5).

**Conclusion:** This study presents the first comprehensive radiobiological evaluation of a laser-driven VHEE beam across multiple biological models. Under the investigated conditions, VHEE and CIEE irradiations exhibit similar biological toxicity. These findings support the feasibility and potential of VHEE generated with LPA for future clinical applications.

## 1. Introduction

The use of Very High Energy Electrons (VHEE) for radiotherapy was first proposed by Des Rosiers et al. [1], who demonstrated, via Monte Carlo simulations, their potential to deliver doses to deep-seated tumors with improved conformality. Unlike conventional electron beams (5–25 MeV), which are limited to superficial treatments [2], VHEE beams (50-300 MeV) can penetrate deeply, enabling treatment of a wider range of malignancies. They also exhibit a sharper lateral penumbra than conventional electrons, and even photons at shallow depths, and may enable uniform dose distributions at depth when delivered in opposed-beam configurations [1, 3]. These early simulations further suggested that VHEE beams are less sensitive to tissue inhomogeneities compared to ion beams and photons, a characteristic later confirmed experimentally by Lagzda et al. [4]. This makes them particularly attractive for treating tumors in heterogeneous regions such as the lungs, bowel, or cervix. Recent treatment planning studies further support their clinical potential, with VHEE treatment plans achieving dose distributions comparable or superior to volumetric modulated arc therapy (VMAT) in pediatric, lung, and prostate cancer cases [5]. From a technological perspective, VHEE beams can be electromagnetically focused and steered with quadrupole magnets, allowing dynamic control of beam trajectory and dose distribution [6, 7]. This capability allows for rapid, flexible, multi-directional dose delivery, streamlining workflow compared to conventional photon-based systems [8]. Despite their promising clinical potential, radiobiological characterization of VHEE beams remains limited. Most experimental studies to date have focused on *in vitro* assessments of relative biological effectiveness (RBE) [9, 10, 11], conducted using radiofrequency (RF) linear accelerators (LINACs), which remain the standard technology for VHEE generating [12, 13, 14]. However, these RF-based systems require large footprint and substantial shielding, and are rarely integrated with biological laboratories, thus complicating routine radiobiological studies.

To overcome these limitations, laser-plasma accelerators (LPAs) have emerged as a promising alternative for VHEE generation [15]. By sustaining gigavolt-per-meter accelerating gradients over millimeter-scale distances, LPAs enable the development of more compact systems that simplify radioprotection requirements and facilitate potential integration into hospital-scale environments. Their compactness further enables novel delivery concepts, including mobile or robotic platforms. For example, Nakajima et al. [16] proposed a robotic, compact gantry-mounted LPA system capable of delivering 50–250 MeV electron beams for imageguided, intensity-modulated radiation therapy (IMRT). Several studies have also explored the feasibility of LPA-based radiotherapy from a dosimetric perspective, including Monte Carlo simulations of dose deposition [17, 18, 19]. Nonetheless, while LPAs offer promising solutions to the spatial and technical constraints of RF systems, most current facilities are still tailored for fundamental physics and lack the biological infrastructure needed for routine radiobiological studies. As a result, only a few radiobiological investigations involving laser-driven VHEE beams have been reported to date [20, 21, 22].

In this study, we report the first comprehensive multi-model investigation of laser-driven VHEE effects across *in vitro, ex vivo*, and *in vivo* assays, using an LPA system adapted for radiobiological research. End-points were selected to probe healthy tissue response at different levels: cell survival in human fibroblasts, proliferation in organotypic mouse lung slices, and developmental toxicity in zebrafish embryos. To ensure clinical relevance and allow benchmarking, all experiments were compared with a conventional 7 MeV LINAC producing what we define as Conventional Intermediate Energy Electrons (CIEE). This study thus establishes a novel preclinical framework for evaluating the radiobiological impact of emerging laser-driven VHEE sources in comparison with clinically established electron modalities.

## 2. Materials and Methods

### 2.1 Ethics Statement

All experimental protocols involving animals complied with French and European regulations on animal care and use (French Decree 2013–118 and EC Directive 2010/63/EU). Zebrafish were obtained from the Laboratory for Optics and Biosciences (Palaiseau, France) and handled in accordance with ethical guidelines. Female C57BL/6J mice (6–10 weeks old) were obtained from Charles River Laboratories (Lyon, France) and housed at the Institut Curie’s animal facility.

### 2.2 Beam Generation and Irradiation Setup

The VHEE beam was generated using the Salle Jaune laser system, a 60 TW Ti:Sapphire laser operating at the Laboratoire d’Optique Appliquée (LOA), Palaiseau, France. The system delivered 30 fs (full width at half maximum) laser pulses with energies exceeding 1.5 J at the focal spot, at a repetition rate of 0.5 Hz. Electron acceleration was achieved inside a vacuum experimental chamber by focusing the laser onto a gas jet of helium (He) mixed with 2% nitrogen (N_2_), expelled from a 4 mm × 250 µm rectangular nozzle (Figure 1a). This laser–plasma interaction generated the VHEE beam, which exited the vacuum chamber through a 1.5 mm thick aluminum (Al) flange and propagates through air towards the biological target. At this position, the broadband electron beam spanned from 50 to 300 MeV. Further details on the beam spectrum are provided in Supplementary Figure A1. Dedicated sample holders (Figures 1b and 1c) were designed to ensure consistent positioning and alignment of the biological targets. The number of laser pulses, and consequently the number of electron shots, was adjusted according to the prescribed dose, while the mean dose rate was maintained at *≈* 24 Gy/min at 0.5 Hz across all experiments.

**Figure 1.**
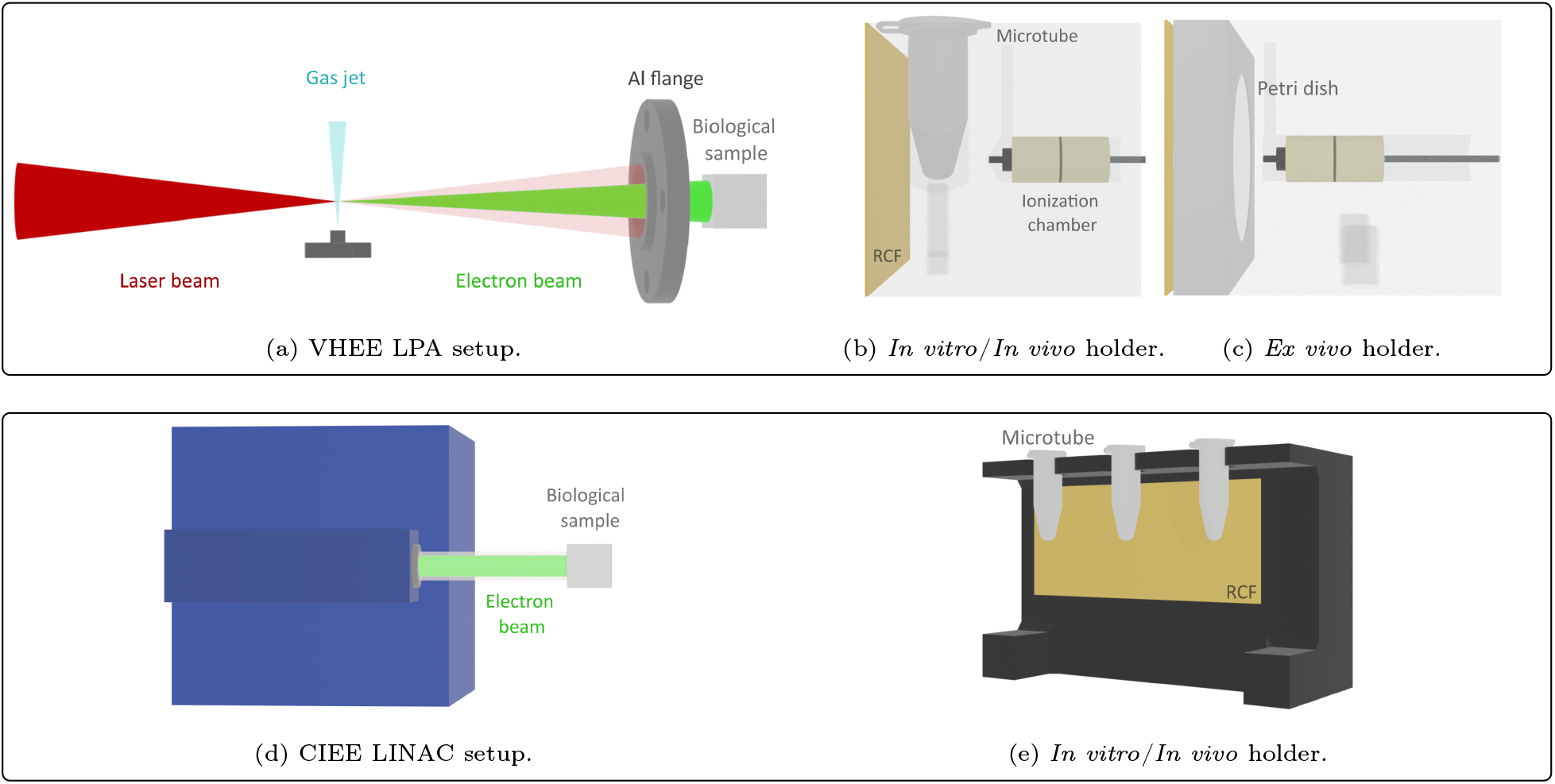
Experimental setups and sample holders used for biological irradiations.Top row (a–c): Laser-plasma–accelerated VHEE setup. (a) Schematic of the LPA system: the laser is focused onto He + 2% N_2_ gas jet, generating the VHEE beam (green). Electrons exit the vacuum chamber through a 1.5 mm thick Al flange toward the biological target. (b) *In vitro*/*in vivo* holder with microtube positioned between a RCF and a ionization chamber for dosimetric verification. (c) *Ex vivo* holder with Petri dish support, RCF, and ionization chamber. Bottom row (d–e): LINAC-based CIEE setup. (d) Schematic of the conventional 7 MeV LINAC. (e) *In vitro*/*in vivo* holder with microtubes placed in front of a RCF, used for dosimetric assessment.

The CIEE beam was generated using the ElectronFLASH LINAC (SIT S.p.A., RD Dept., Roma, Italy) at Institut Curie. The electron energy was set to 7 MeV, and samples were irradiated at a source-to-surface distance of 1.1 m, ensuring uniform dose delivery across the irradiated field. A fixed pulse repetition rate of 10 Hz was used, delivering a mean dose rate of approximately ≈ 30 Gy/min for all experiments. The LINAC system and the customized biological sample holder used for both *in vitro* and *in vivo* irradiations are illustrated in Figures 1d and 1e. The *ex vivo* samples were irradiated using standard culture plates.

### 2.3 Dosimetry

For VHEE irradiations, a Razor Nano Chamber (IBA Dosimetry, Schwarzenbruck, Germany) was used for real-time dose monitoring. Prior to each experiment, a calibration was performed to determine the dose ratio between the sample position and the chamber’s placement during irradiation. This allowed estimation of the dose-per-pulse and calculation of the required number of shots to reach the prescribed dose. Additionally, 1 cm water-equivalent material bricks were placed in front of the sample to adjust the beam size and average dose rate. Real-time readings from the chamber were used to monitor dose delivery and ensure accurate irradiation of the target.

For CIEE irradiations, the LINAC’s internal monitoring ionization chamber was calibrated to relate Monitor Units (MU) to absolute dose. The LINAC was programmed to automatically stop beam delivery once the desired dose was reached, based on the cumulative MUs.

### 2.4 In Vitro Cell Survival Assay

The immortalized MRC5-hTERT human fibroblast cell line was cultured in Dulbecco’s Modified Eagle Medium (DMEM + GlutaMAX, Gibco), supplemented with 10% fetal calf serum (PAA) and Pen-Strep antibiotics (10 U/mL penicillin, 10 µg/mL streptomycin, Gibco), as described in the established protocol by Bayart et al. [23]. Cells were maintained as monolayers at 37°C in a humidified atmosphere with 5% CO_2_. For irradiation, 10^4^ cells were transferred into Eppendorf tubes (2 mL for VHEE, 0.5 mL for CIEE) containing growth medium, centrifuged, and irradiated at room temperature (22–24°C). Irradiations were performed across three days. Post-exposure, each cell pellet was resuspended and evenly seeded into four wells of a 12-well plate. Fresh medium was added to each well to reach a final volume of 2 mL. After incubation for five generations (six days), cells were harvested using 0.2 mL of Accutase (EMD Millipore) and neutralized with an equal volume of medium. Viable cells were counted using an ORFLO Moxi Mini Automated Cell Counter (Type S cassette). Dose–response survival curves were established for both VHEE and CIEE conditions, with doses ranging from 2 to 10 Gy.

### 2.5 Mouse PCLS Preparation and Cell Division Assay

Organotypic precision-cut lung slices (PCLS) were prepared following the protocol described by Dubail et al. [24]. Lungs from anesthetized C57BL/6J mice (6–10 weeks old) were sectioned into 300 µm slices using a vibratome and maintained in culture at 37°C with 5% CO_2_ until irradiation. For VHEE irradiation, five slices were randomly assigned to each dose groups (3, 6, and 9 Gy) and irradiated individually in Petri dishes. Irradiations were carried out over three independent days, using one mouse per day. Cell proliferation was assessed 24 hours post-irradiation using a Click-iT™ chemistry protocol based on the incorporation of 5-ethynyl-2’-deoxyuridine (EdU) (BCK-EdUPro-FC647). Imaging was performed using a Nikon Spinning Disk TIRF-FRAP microscope, with approximately 5-7 fields of view (FOVs) acquired per slice and analysed with IMARIS (Bitplane). CIEE irradiations were conducted using the same lung slice preparation, dose grouping, and proliferation assessment protocol.

### 2.6 Zebrafish Embryo Irradiation and Morphological Quantification

Wild-type AB zebrafish embryos were maintained in E3 medium at 28°C until irradiation. At 5 hours post-fertilization, embryos were transferred into Eppendorf tubes (2 mL for VHEE, 0.5 mL for CIEE) filled with E3 medium and irradiated at room temperature (22–24°C) with either VHEE or CIEE beams, at doses of 6 and 9 Gy. Each dose group under both irradiation modalities was independently replicated across three experimental days. For each target dose, approximately 111 embryos were irradiated with VHEE and around 70 with CIEE. At 5 days post-irradiation, embryos were fixed in 10% formalin and imaged using an Echo Rebel microscope (4× objective). Morphological abnormalities were quantitatively assessed, focusing on body length and spinal curvature. Both parameters were computed using a custom Python-based analysis tool with manual midline tracing via a graphical interface. Body length was calculated as the distance along the vertebral axis from the anterior tip of the head to the end of the vertebral column, excluding the caudal fin fold. Spinal curvature was assessed using two complementary metrics derived from a separately traced midline along the tail region: the mean local curvature (average angular deviation between adjacent triplets of points) and the maximum angle curvature (largest angular deviation along the spine).

### 2.7 Statistical Analysis

For cell survival, lung slice, and zebrafish assays, irradiated sample values were normalized to the corresponding non-irradiated (NI) control from the same experimental day and averaged across biological replicates. Dose–response curves for cell survival were fitted using the linear–quadratic model in GraphPad Prism (version 10.3.1). Data distributions from lung slice and zebrafish assays were visualized using violin plots, with overlaid boxplots showing the mean, median, interquartile range (IQR), and whiskers extending to 1.5× IQR. Statistical comparisons for lung slice and zebrafish data were performed using the Kruskal–Wallis test to assess differences across groups, followed by Dunn’s post hoc test with Holm correction for multiple comparisons (scipy.stats and scikit-posthocs, Python v3.X, SciPy v1.11.4). A two-sided p-value <0.05 was considered statistically significant.

## 1. Results

### 3.1 Human Fibroblasts Exhibit Comparable Radiosensitivity to Laser-Driven VHEE and CIEE

As no prior radiobiological experiments had been conducted using the Salle Jaune laser facility, we first assessed cell survival in MRC-hTERT human fibroblasts irradiated with the laser-driven VHEE beam. Figure 2 shows the dose-response curves for both VHEE and CIEE irradiations, fitted using the linear-quadratic model (R^2^ = 0.98 for VHEE; R^2^ = 0.97 for CIEE). The dose required to reduce cell survival to 10% (D_10_) was 6.71 ± 1.09 Gy for VHEE and 6.38 ± 0.42 Gy for CIEE (mean ± standard error of the mean, SEM), with no statistically significant difference between the two (p = 0.70). This result indicates that VHEE and CIEE exhibit comparable radiosensitivity in this healthy cell model, further supported by the similarity in curve shapes and overlapping dose uncertainties.

**Figure 2.**
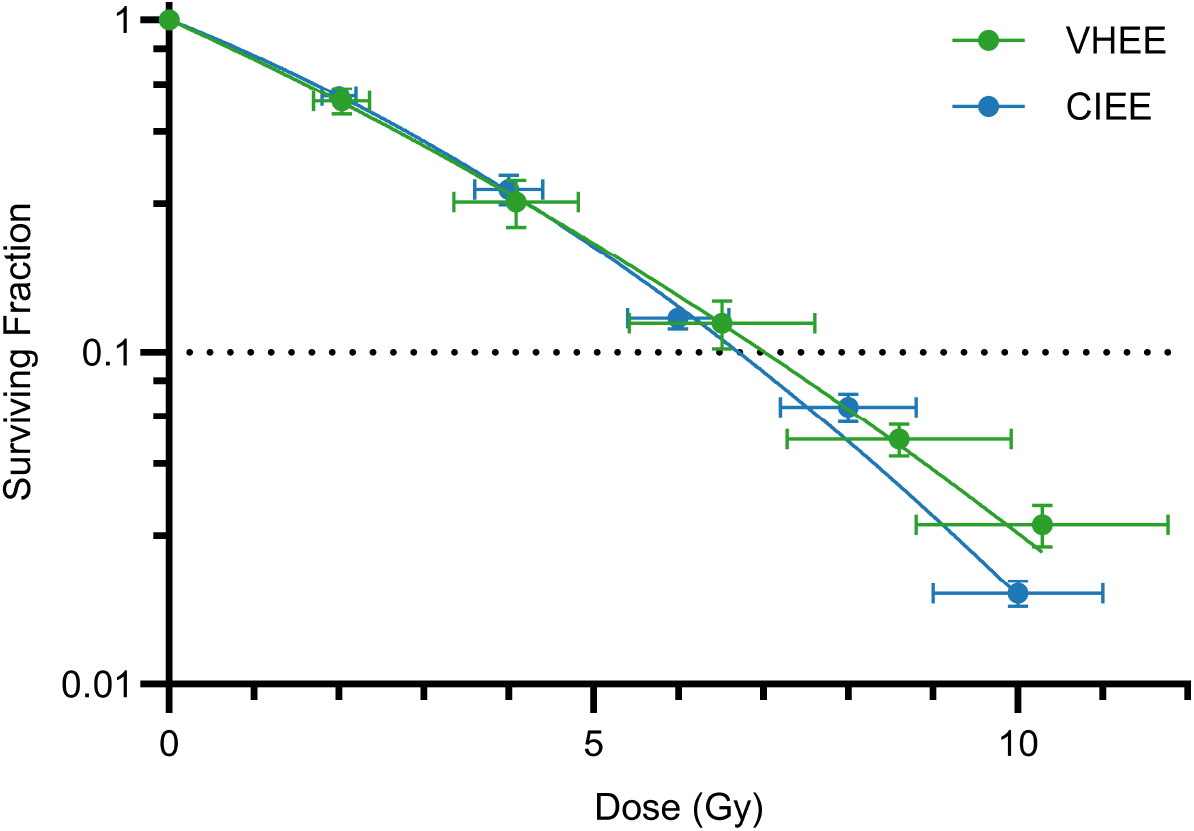
Dose–response survival curves for MRC5-hTERT human fibroblasts irradiated with either laser-driven VHEE (green) or CIEE (blue). Each point represents the mean surviving fraction ± SEM, with horizontal error bars indicating dose uncertainty. Curves were fitted using the linear–quadratic model (R^2^ > 0.93 for VHEE, R^2^ > 0.97 for CIEE). The dotted line marks the D_10_, which was not significantly different between the two modalities (p = 0.70).

### 3.2 PCLS Reveal Similar Cell Division Decrease After VHEE and CIEE Exposure

To investigate beam toxicity in a tissue-relevant model, we used mouse PCLS, which preserve tissue architecture and much of the lung microenvironment. EdU^+^ cell quantification following exposure to target doses of 3, 6, and 9 Gy revealed a progressive, dose-dependent reduction in cell proliferation for both irradiation modalities (Figure 3). Relative to NI controls (100%), VHEE exposure reduced proliferation to 48.5% ± 2.1, 31.6% ± 2.3, and 17.0% ± 1.5 (mean ± SEM) at measured doses of 3.07 ± 0.65, 5.68 ± 1.27, and 8.73 ± 2.32 Gy, respectively. CIEE yielded comparable reductions: 46.9% ± 2.7, 32.2% ± 2.4, and 20.0% ± 2.3 (mean ± SEM) at 3.00 ± 0.12, 6.00 ± 0.24, and 9.00 ± 0.36 Gy, respectively. Inter-dose comparisons within each modality confirmed statistically significant reductions in proliferation (VHEE: p *≤* 1.88 *×* 10^−3^; CIEE: p *≤* 2.48 *×* 10^−2^). However, no significant differences were observed between VHEE and CIEE at matched dose levels, supporting their comparable biological effects in healthy lung tissue. Complete data are reported in Supplementary Tables A1 and A2.

**Figure 3.**
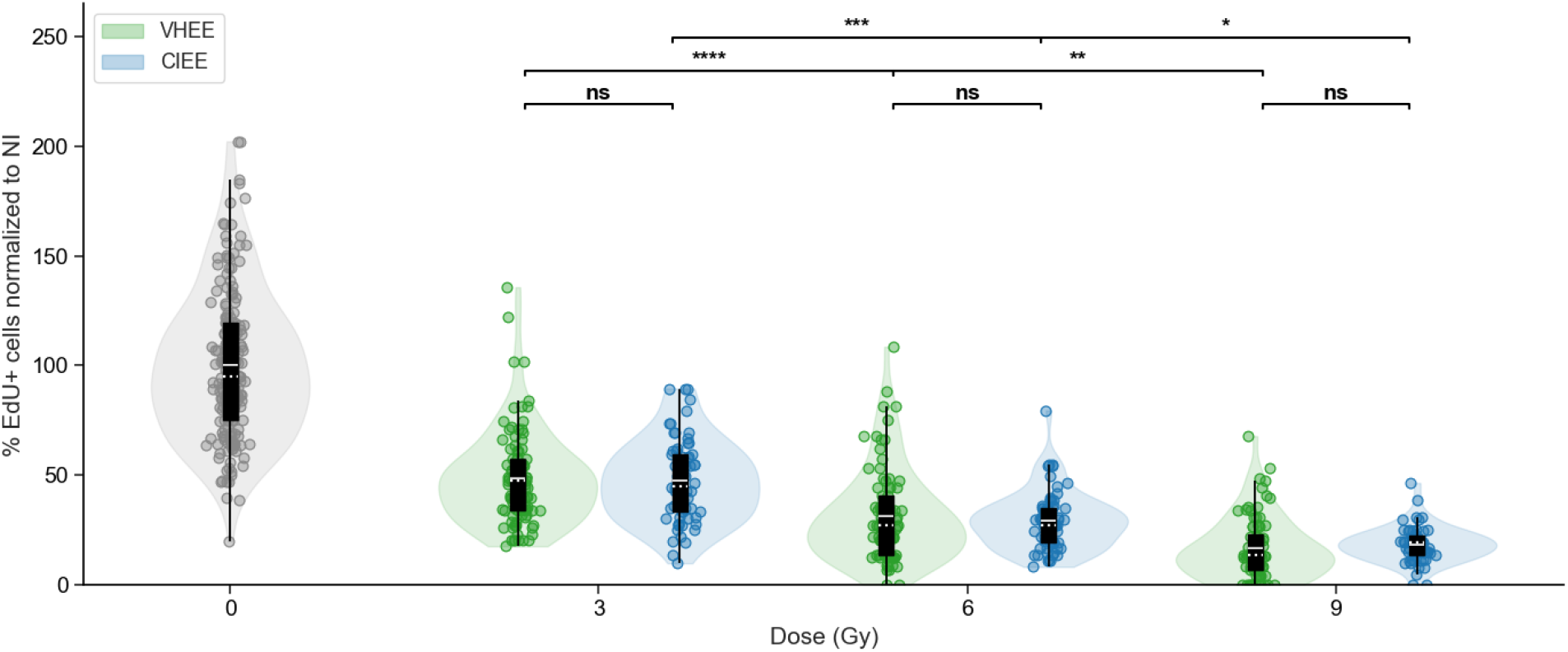
Quantification of the proportion of EdU^+^ cells in PCLS, normalized to NI controls, after radiation exposure to VHEE (green) or CIEE (blue) at target doses of 3, 6, and 9 Gy. Violin plots show the distribution of FOV measurements; overlaid boxplots (mean: solid white line; median: dotted white line; box: IQR; whiskers: 1.5× IQR). Statistical comparisons were performed using the Kruskal–Wallis test followed by Dunn’s post hoc test with Holm correction. Significance leves: ns, not significant; * p-value <0.05; ** p-value <0.01; *** p-value <0.001; **** p-value <0.0001.

### 3.3 Zebrafish Embryos Show Comparable Dose-Dependent Toxicity from VHEE and CIEE

Zebrafish embryos were used as an *in vivo* model to assess developmental toxicity following whole-body exposure to either VHEE or CIEE irradiation. Morphological analyses at 5 days post-irradiation revealed a dose-dependent effect across all evaluated metrics: normalized body length, normalized mean local curvature, and normalized maximum angle curvature (Figures 4b, 4c and 4d). A progressive reduction in normalized body length was observed with increasing dose for both modalities. At 6 Gy (actual dose: 6.3 ± 0.7 Gy VHEE, 6.0 ± 0.24 Gy CIEE), reductions relative to NI controls were modest (−3.3% VHEE and -2.7% CIEE), while at 9 Gy (actual dose: 9.5 ± 0.9 Gy VHEE, 9 ± 0.36 Gy CIEE), shortening became more pronounced (−13.4% VHEE and -11.8% CIEE). Curvature metrics increased with dose: at 9 Gy, mean local curvature reached 4.2 ± 0.5 (VHEE) and 4.1 ± 0.5 (CIEE), while maximum angle curvature rose to 6.0 ± 0.8 (VHEE) and 6.3 ± 1.0 (CIEE). Statistical analysis confirmed significant intra-modality effects between 6 and 9 Gy for all metrics (VHEE: p*≤* 2.14 *×* 10^−4^; CIEE: p≤ 2.52 × 10^−5^). Representative images of embryos exposed to 0 Gy and 9 Gy (Figure 4a) illustrate the curvature enhancement induced by high-dose exposure. In contrast, no statistically significant inter-modality differences were observed at either 6 or 9 Gy (p≥ 0.5), supporting comparable morphological outcomes for VHEE and CIEE. Complete data are reported in Supplementary Tables A3 and A4.

**Figure 4.**
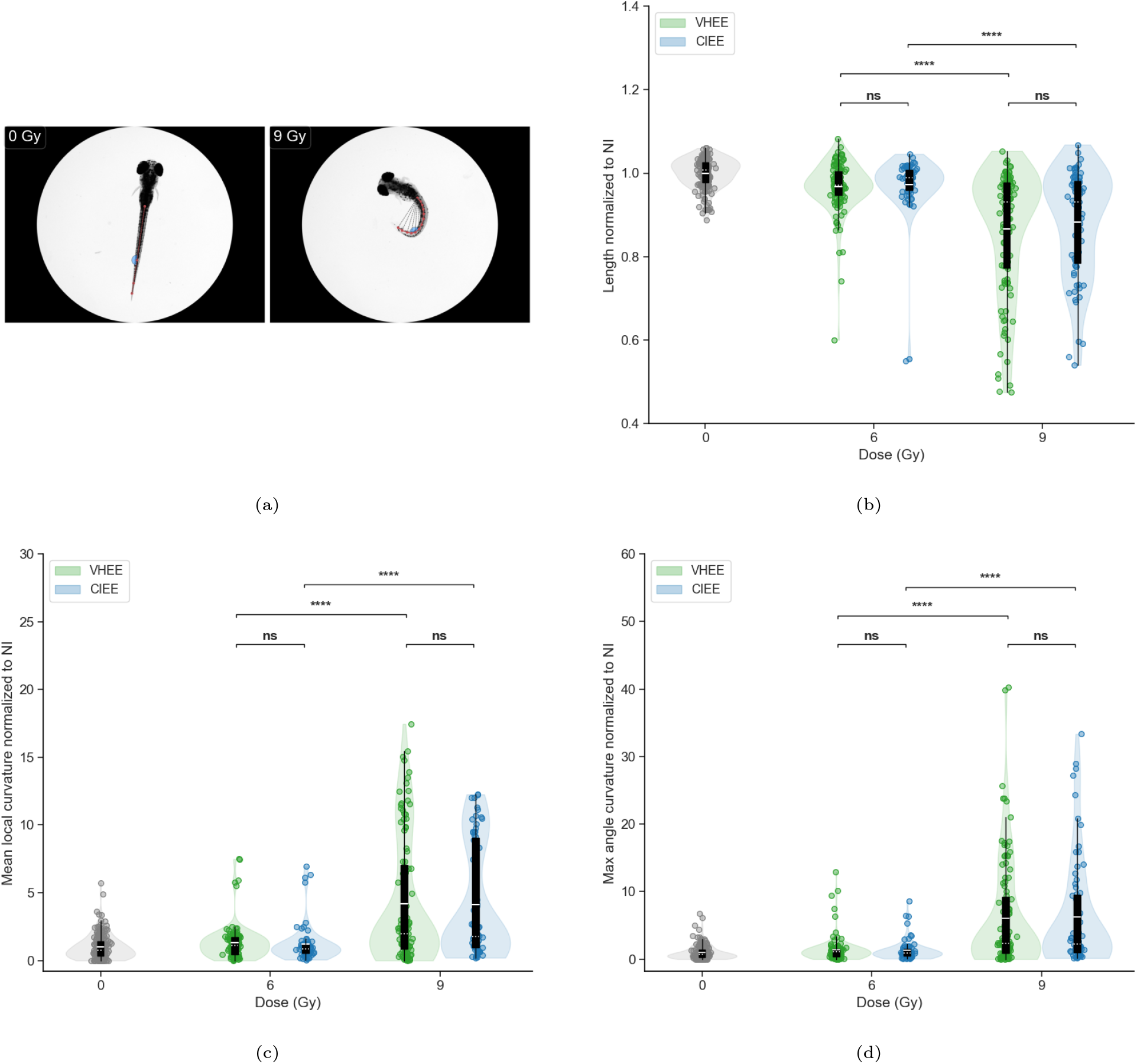
Morphological effects of VHEE and CIEE irradiation on zebrafish embryos at 5 days post-irradiation. (a) Representative images of zebrafish at 5 days post-irradiation with 0 Gy (NI) and 9 Gy VHEE, showing increased spinal curvature (curvature traces in red, maximum curvature angle in blue). (b) Normalized body length at 0, 6, and 9 Gy for VHEE (green) and CIEE (blue). (c) Normalized mean local curvature. (d) Normalized maximum angle curvature. Violin plots show full data distributions with overlaid boxplots (mean: solid white line; median: dotted white line; box: IQR; whiskers: 1.5× IQR). Each dot represents a single embryo. Statistical comparisons: Kruskal–Wallis test with Dunn’s post hoc and Holm correction. Significance levels: ns, not significant; * p-value <0.05; ** p-value <0.01; *** p-value <0.001; **** p-value <0.0001.

## 4. Discussion

VHEE are emerging as a promising modality for radiotherapy, offering deep tissue penetration and sharp lateral dose gradients. These characteristics make them particularly well-suited for treating tumors in anatomically complex or heterogeneous regions, where precision and robustness against tissue inhomogeneities are critical. When combined with LPA technology, VHEE delivery becomes more feasible in hospital-scale environments, as millimeter-scale acceleration enables compact systems with reduced radioprotection requirements.

In this study, we present the first comprehensive radiobiological evaluation of laser-accelerated VHEE beams across *in vitro, ex vivo*, and *in vivo* models. The successful adaptation of our LPA system for biological irradiation enabled stable and reproducible beam delivery under controlled conditions, a prerequisite to translate the use of LPAs into radiobiology applications.

Our results consistently show that VHEE irradiation induces biological responses comparable to those observed with a conventional 7 MeV electron beam (CIEE). In human fibroblasts, dose–response curves revealed minimal differences in D_10_ values between the two modalities, demonstrating both the feasibility of conducting radiobiological assays with our laser-driven VHEE system and their comparable toxicity in this cell model. This fundamental result motivated the extension of our investigation to more complex systems. *Ex vivo*, proliferation assays in organotypic mouse PCLS revealed a dose-dependent reduction in EdU^+^ cells, with no statistically significant differences between the two beams at any dose. *In vivo*, zebrafish embryos exhibited comparable morphological changes following VHEE or CIEE exposure, including dose-dependent effects on body length, mean local curvature, and maximum angle curvature. Together, these results indicate that laser-driven VHEE exhibit toxicity comparable to that of CIEE across models of increasing biological complexity.

While our study focused on healthy tissue models and short-term endpoints, this approach provided a robust and controlled framework to compare the biological effects of laser-driven VHEE with those of a conventionally accelerated 7 MeV electron beam. In the systems, fibroblast monolayers, lung slices, and zebrafish embryos, electron penetration depth was sufficient to ensure full irradiation, which may explain the close agreement observed between the two modalities. However, one of the key advantages of VHEE is their ability to penetrate deeply into tissue. To fully explore this potential, future studies should involve anatomically larger systems, in which tissue thickness becomes a limiting factor for low-energy electron beams, allowing the deep-penetrating properties of VHEE to be more meaningfully evaluated, particularly in terms of spatial dose distribution and Bragg-peak–like deposition enabled by quadrupole focusing [1, 6, 7]. Beyond spatial control, a remarkable feature of LPA-based systems lies in their temporal structure. Our laser-based electron accelerator delivers dose in sub-picosecond bursts with extremely high instantaneous peak dose rates (exceeding 10^9^ Gy*/*s). Although current dose-per-pulse and repetition rate constraints preclude FLASH delivery [25], ongoing technological progress may soon enable FLASH-compatible LPA-driven VHEE. Even before achieving FLASH condition, the ultra-short pulsed nature of laser-based systems allows exploration of fast dose fractionation irradiation modality, by tuning the temporal delay between pulses. Recent work with laser-accelerated protons has shown that modifying pulse timing can impair DNA repair mechanisms, leading to enhanced tumor cell mortality and potentially improved therapeutic indices [23]. Similar radiobiological effects could be explored using laser-accelerated electrons, enabling novel strategies for optimizing temporal delivery in radiotherapy. Looking ahead, preclinical studies incorporating tumor models, long-term toxicity outcomes (e.g., fibrosis, regeneration), and direct comparisons between RF- and laser-driven VHEE platforms will be essential to advancing clinical translation.

## 5. Conclusion

This study presents the first comprehensive radiobiological evaluation of laser-driven VHEE across *in vitro, ex vivo*, and *in vivo* models. Through the successful adaptation of our LPA system, we achieved stable and reproducible irradiation across increasing levels of biological complexity, from fibroblast survival to tissue-level proliferation and whole-organism morphological alterations. Across all systems, VHEE-induced toxicity was comparable to that of CIEE beams. These findings provide a robust foundation for future preclinical studies with VHEE beams, particularly when delivered via LPA-based systems, which offer unique advantages in compactness, flexibility, and ultrashort beam temporal structure. While this work focused on healthy tissues and conventional dose rates, it provides a valuable reference for future investigations into tumor-bearing models, long-term toxicity outcomes, and advanced temporal delivery strategies such as FLASH and fast fractionation. By bridging cutting-edge accelerator physics with rigorous biological testing, this study positions laser-driven VHEE as a novel and promising tool in the evolving landscape of particle therapy.

## Notes

### Competing Interest Statement

The authors have declared no competing interest.

### Summary of Updates

This version of the manuscript includes updated and more detailed information regarding the energy spectrum of the electron beam produced by the laser-plasma accelerator. Experimental and simulated spectra have been added in the Supplementary Materials to better characterize the actual irradiation conditions.

### Funding

- CNRS: MITI/BioRapide
- Île-de-France: Sesame/IDRA
- ANR: FemtoDose
- FondationX: BioRapideHC
- ANR/France 2030: ANR-24-FRAT-001
- INSERM: 19CR057-00

