## Supplementary Figure A1, Supplementary Tables A1, A2, A3, and A4 for "Multiscale radiobiological assessment of laser-driven very high energy electrons versus conventional electrons"

### SUPPLEMENTARY MATERIALS

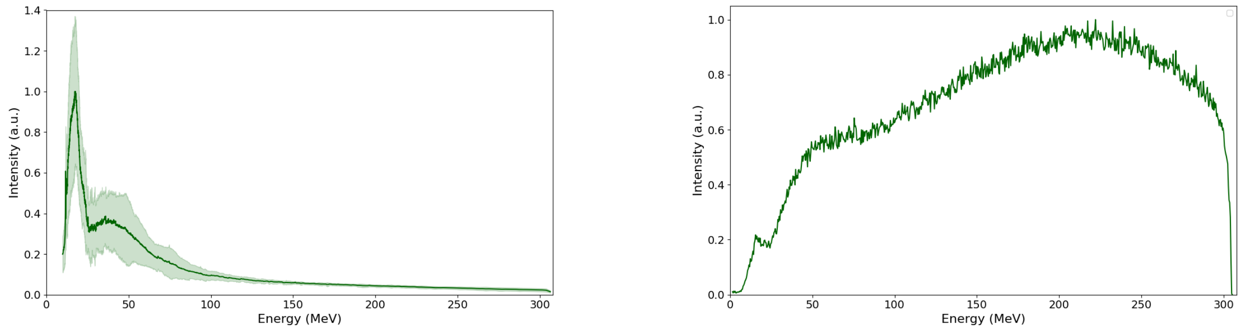


(a) (b)

Figure A1: (a) Experimental average spectrum of the laser-driven VHEE beam over 60 shots, measured at the laser–plasma interaction point. The shaded area represents the standard deviation. (b) Simulated Monte Carlo spectrum (GEANT4) at the biological sample location. The experimental VHEE spectrum measured at the source was used as input. The beamline geometry included a 1.5 mm aluminum vacuum–air flange with an attached 1 mm brass diffuser, followed by 16 cm of air propagation. A total of $1.2 \times 10^7$ events were simulated to ensure statistical reliability.

Table A1: Quantification of EdU$^+$ cells in mouse precision-cut lung slices (PCLS) following irradiation with VHEE or CIEE at different target doses. Measured doses are reported as mean ± SD. The proportion of EdU$^+$ cells is expressed as percentage relative to non-irradiated (NI) controls and reported as mean ± SEM.

| **Modality** | **Target Dose (Gy)** | **Measured Dose (Gy) (mean ± SD)** | **EdU$^+$ cells (% of NI) (mean ± SEM)** |
|---|---|---|---|
| CIEE | 0 | – | 100.00 ± 3.94 |
| | 3 | 3.000 ± 0.12 | 47.62 ± 2.23 |
| | 6 | 6.000 ± 0.24 | 29.27 ± 1.57 |
| | 9 | 9.000 ± 0.36 | 18.45 ± 1.05 |
| VHEE | 0 | – | 100.00 ± 3.18 |
| | 3 | 3.072 ± 0.65 | 48.49 ± 2.09 |
| | 6 | 5.680 ± 1.27 | 31.55 ± 2.28 |
| | 9 | 8.734 ± 2.32 | 17.01 ± 1.53 |

Table A2: Statistical analysis of $EdU^{+}$ cell proportions in mouse PCLS across all irradiation conditions. A global comparison using the Kruskal–Wallis test ($H = 450.900$, $p = 2.845 \times 10^{-93}$) was followed by Dunn's post hoc test with Holm correction for pairwise comparisons. Significance levels are indicated as follows: $p < 0.05$ (*), $p < 0.01$ (**), $p < 0.001$ (***), and $p < 0.0001$ (****). Comparisons not reaching statistical significance are indicated as (ns).

| **Condition A** | **Condition B** | **p-value** |
|---|---|---|
| 0 Gy VHEE | 0 Gy CIEE | $1.00 \times 10^{0}$ (ns) |
| 0 Gy VHEE | 3 Gy CIEE | $3.72 \times 10^{-10}$ (****) |
| 0 Gy VHEE | 6 Gy CIEE | $4.22 \times 10^{-27}$ (****) |
| 0 Gy VHEE | 9 Gy CIEE | $1.00 \times 10^{-41}$ (****) |
| 0 Gy VHEE | 3 Gy VHEE | $2.25 \times 10^{-12}$ (****) |
| 0 Gy VHEE | 6 Gy VHEE | $2.00 \times 10^{-30}$ (****) |
| 0 Gy VHEE | 9 Gy VHEE | $1.30 \times 10^{-53}$ (****) |
| 3 Gy VHEE | 3 Gy CIEE | $1.00 \times 10^{0}$ (ns) |
| 3 Gy VHEE | 6 Gy CIEE | $1.50 \times 10^{-4}$ (***) |
| 3 Gy VHEE | 9 Gy CIEE | $7.46 \times 10^{-12}$ (****) |
| 3 Gy VHEE | 6 Gy VHEE | $8.09 \times 10^{-5}$ (****) |
| 3 Gy VHEE | 9 Gy VHEE | $2.04 \times 10^{-15}$ (****) |
| 6 Gy VHEE | 6 Gy CIEE | $1.00 \times 10^{0}$ (ns) |
| 6 Gy VHEE | 9 Gy CIEE | $1.21 \times 10^{-2}$ (*) |
| 6 Gy VHEE | 9 Gy VHEE | $1.88 \times 10^{-3}$ (**) |
| 9 Gy VHEE | 9 Gy CIEE | $1.00 \times 10^{0}$ (ns) |
| 3 Gy VHEE | 0 Gy CIEE | $2.86 \times 10^{-9}$ (****) |
| 6 Gy VHEE | 0 Gy CIEE | $8.61 \times 10^{-23}$ (****) |
| 9 Gy VHEE | 0 Gy CIEE | $5.08 \times 10^{-40}$ (****) |
| 6 Gy VHEE | 3 Gy CIEE | $6.26 \times 10^{-4}$ (***) |
| 9 Gy VHEE | 3 Gy CIEE | $1.71 \times 10^{-12}$ (****) |
| 9 Gy VHEE | 6 Gy CIEE | $6.19 \times 10^{-3}$ (**) |

Table A3: Quantification of zebrafish morphological metrics following irradiation with VHEE or CIEE at different target doses. Measured doses are reported as mean ± SD. Morphological metrics (normalized length, mean local curvature, and maximum angle curvature) are normalized to NI controls and expressed as mean ± SEM.

| **Metric** | **Modality** | **Target Dose (Gy)** | **Measured Dose (Gy) (mean ± SD)** | **Computed Metric (mean ± SEM)** |
|---|---|---|---|---|
| Normalized Length | CIEE | 0 | – | 1.000 ± 0.004 |
| | VHEE | 0 | – | 1.000 ± 0.003 |
| | CIEE | 6 | 6.00 ± 0.24 | 0.973 ± 0.009 |
| | VHEE | 6 | 6.33 ± 0.68 | 0.967 ± 0.006 |
| | CIEE | 9 | 9.00 ± 0.36 | 0.882 ± 0.015 |
| | VHEE | 9 | 9.45 ± 0.90 | 0.866 ± 0.014 |
| Normalized Mean Local Curvature | CIEE | 0 | – | 1.000 ± 0.082 |
| | VHEE | 0 | – | 1.000 ± 0.097 |
| | CIEE | 6 | 6.00 ± 0.24 | 1.224 ± 0.165 |
| | VHEE | 6 | 6.33 ± 0.68 | 1.311 ± 0.119 |
| | CIEE | 9 | 9.00 ± 0.36 | 4.114 ± 0.494 |
| | VHEE | 9 | 9.45 ± 0.90 | 4.195 ± 0.449 |
| Normalized Max Angle Curvature | CIEE | 0 | – | 1.000 ± 0.103 |
| | VHEE | 0 | – | 1.000 ± 0.111 |
| | CIEE | 6 | 6.00 ± 0.24 | 1.359 ± 0.184 |
| | VHEE | 6 | 6.33 ± 0.68 | 1.417 ± 0.177 |
| | CIEE | 9 | 9.00 ± 0.36 | 6.270 ± 0.956 |
| | VHEE | 9 | 9.45 ± 0.90 | 6.019 ± 0.781 |

Table A4: Pairwise statistical comparisons of zebrafish morphological metrics across all irradiation conditions and doses. A Kruskal–Wallis test was performed for each metric: normalized length (H = 143.72, $p = 2.89 \times 10^{-29}$), mean local curvature (H = 75.27, $p = 8.16 \times 10^{-15}$), and maximum angle curvature (H = 89.09, $p = 1.04 \times 10^{-17}$). These were followed by Dunn's post hoc test with Holm correction for multiple comparisons. Pairwise p-values for each metric are reported in the table. Significance levels are indicated as follows: $p < 0.05$ (*), $p < 0.01$ (**), $p < 0.001$ (***), and $p < 0.0001$ (****). Comparisons not reaching statistical significance are indicated as (ns).

| **Condition A** | **Condition B** | **p-value (Length)** | **p-value (Mean Curvature)** | **p-value (Max Angle Curvature)** |
|---|---|---|---|---|
| 0 Gy VHEE | 0 Gy CIEE | $2.35 \times 10^{-1}$ (ns) | $1.00 \times 10^{0}$ (ns) | $1.00 \times 10^{0}$ (ns) |
| 6 Gy VHEE | 0 Gy VHEE | $1.87 \times 10^{-5}$ (****) | $9.76 \times 10^{-2}$ (ns) | $1.16 \times 10^{-1}$ (ns) |
| 6 Gy VHEE | 0 Gy CIEE | $2.02 \times 10^{-4}$ (***) | $1.00 \times 10^{0}$ (ns) | $1.00 \times 10^{0}$ (ns) |
| 6 Gy VHEE | 6 Gy CIEE | $4.95 \times 10^{-1}$ (ns) | $1.00 \times 10^{0}$ (ns) | $1.00 \times 10^{0}$ (ns) |
| 6 Gy VHEE | 9 Gy VHEE | $1.04 \times 10^{-5}$ (****) | $1.88 \times 10^{-4}$ (***) | $1.41 \times 10^{-5}$ (****) |
| 6 Gy VHEE | 9 Gy CIEE | $8.35 \times 10^{-4}$ (***) | $2.14 \times 10^{-4}$ (***) | $3.04 \times 10^{-6}$ (****) |
| 9 Gy VHEE | 0 Gy VHEE | $2.80 \times 10^{-20}$ (****) | $4.49 \times 10^{-10}$ (****) | $1.20 \times 10^{-11}$ (****) |
| 9 Gy VHEE | 0 Gy CIEE | $1.68 \times 10^{-15}$ (****) | $2.52 \times 10^{-5}$ (****) | $4.59 \times 10^{-6}$ (****) |
| 9 Gy VHEE | 6 Gy CIEE | $2.04 \times 10^{-7}$ (****) | $2.15 \times 10^{-5}$ (****) | $1.92 \times 10^{-4}$ (***) |
| 9 Gy VHEE | 6 Gy VHEE | $1.04 \times 10^{-5}$ (****) | $1.88 \times 10^{-4}$ (***) | $1.41 \times 10^{-5}$ (****) |
| 9 Gy VHEE | 9 Gy CIEE | $1.00 \times 10^{0}$ (ns) | $1.00 \times 10^{0}$ (ns) | $1.00 \times 10^{0}$ (ns) |
| 6 Gy CIEE | 0 Gy CIEE | $4.41 \times 10^{-2}$ (*) | $1.00 \times 10^{0}$ (ns) | $1.00 \times 10^{0}$ (ns) |
| 6 Gy CIEE | 0 Gy VHEE | $3.10 \times 10^{-2}$ (*) | $1.00 \times 10^{0}$ (ns) | $2.35 \times 10^{-1}$ (ns) |
| 9 Gy CIEE | 0 Gy CIEE | $1.30 \times 10^{-11}$ (****) | $2.52 \times 10^{-5}$ (****) | $8.92 \times 10^{-7}$ (****) |
| 9 Gy CIEE | 0 Gy VHEE | $2.71 \times 10^{-14}$ (****) | $3.43 \times 10^{-9}$ (****) | $7.32 \times 10^{-12}$ (****) |
| 9 Gy CIEE | 6 Gy CIEE | $2.58 \times 10^{-5}$ (****) | $2.52 \times 10^{-5}$ (****) | $3.31 \times 10^{-5}$ (****) |
